## Supplementary Figures for "Microbiome meta-analysis and cross-disease comparison enabled by the SIAMCAT machine-learning toolbox"

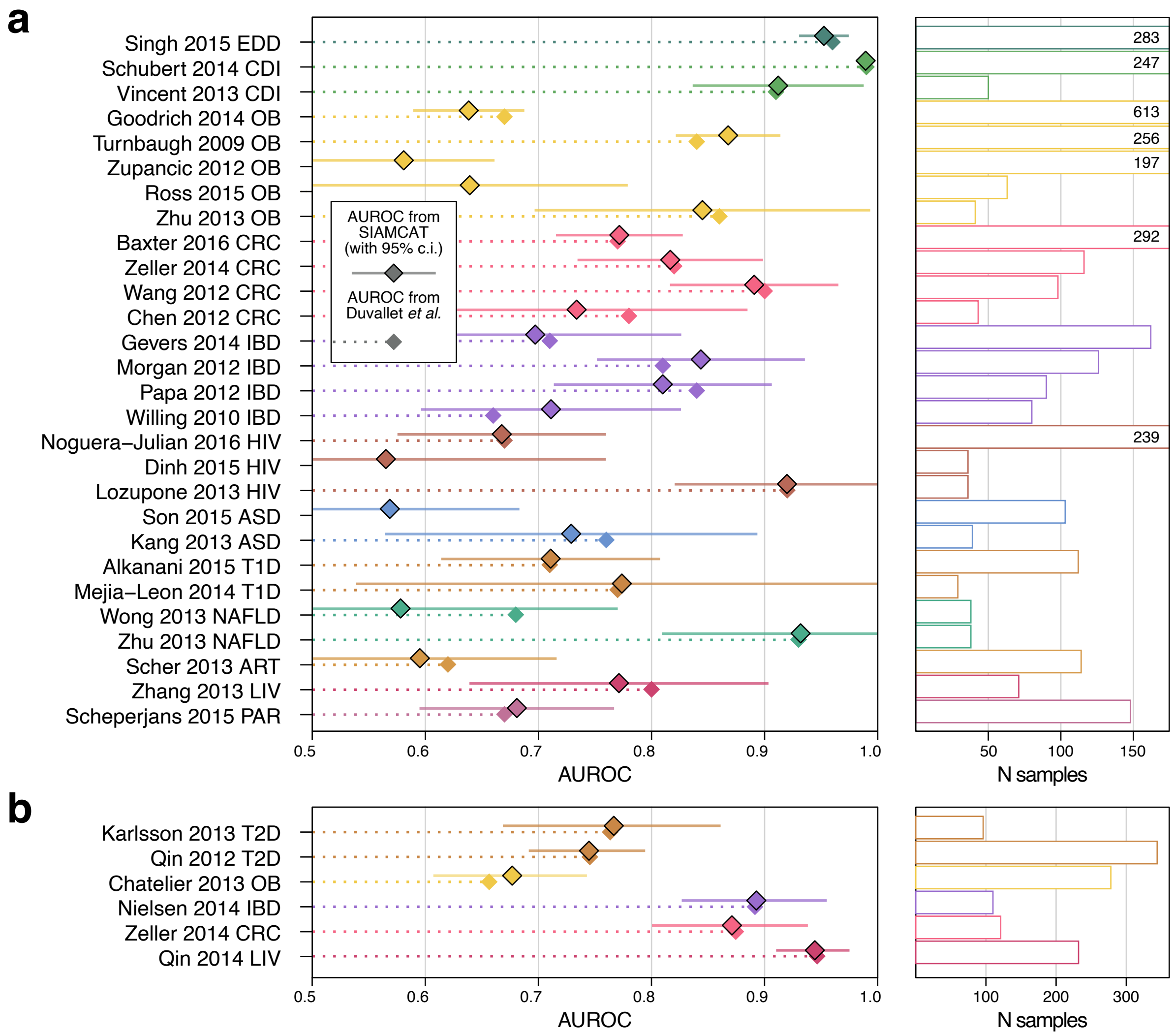


**Supplementary Figure 1: Reproduction of previous machine learning meta-analyses using SIAMCAT**

To show how SIAMCAT can reproduce the results of previous meta-analyses, we reanalyzed the data from Duvallet *et al.* **(a)** and Pasolli *et al.* **(b)** (see references in the main text). SIAMCAT workflows were implemented to fully recapitulate the machine learning workflows as described in the respective publications, using the `randomForest` ML algorithm in both cases. Cross-validation performance quantified by AUROC for discriminating between diseased patients and controls is indicated by diamonds with black borders (95% confidence intervals denoted by horizontal lines) for the SIAMCAT reproduction. The AUROC values reported in the publications are indicated by diamonds without borders. The sample size of each dataset is given as additional panel (cut at N = 200 and given by numbers instead). For all classification tasks, the reported results fall within the confidence interval of the SIAMCAT results.

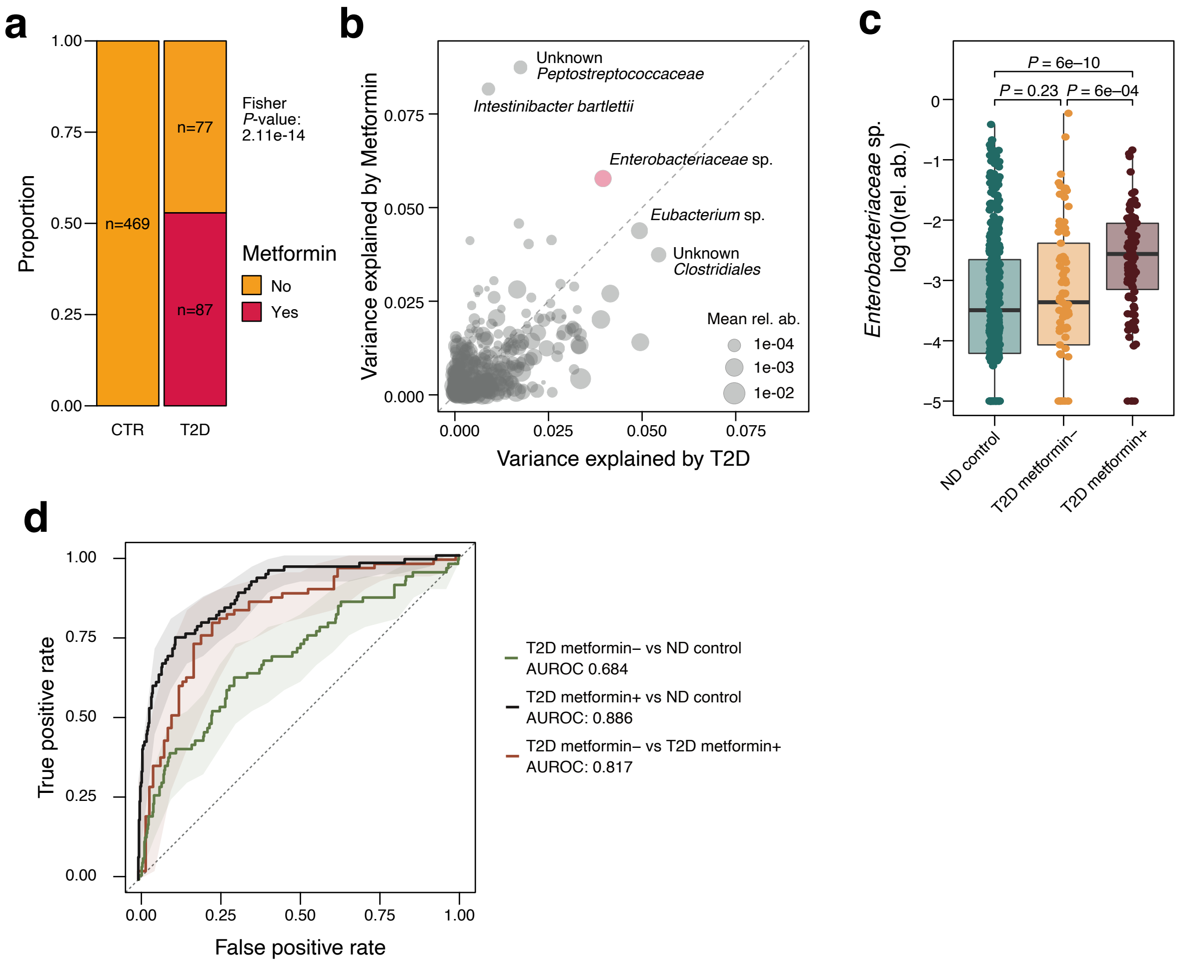


**Supplementary Figure 2: SIAMCAT can detect confounding factors such as metformin treatment**
To show how SIAMCAT can aid confounder detection, we re-analyzed the data from (Forslund *et al.* 2015), which included samples with type 2 diabetes (T2D) and non-diabetic (ND) controls and information about metformin treatment. **(a)** Output of the *check.confounders* function for the data from Forslund *et al.* shows that only T2D cases were treated with metformin, suggesting that metformin-treatment could confound the associations between microbiome features and T2D. **(b)** Analysis of variance (using ranked abundance data) shows that many species differ as much (or more) by metformin treatment as by T2D status. Extreme cases of confounding are highlighted. Dot size is proportional to the mean relative abundance across samples. **(c)** Relative abundance of *Enterobacteriacaea* sp. are significantly larger for metformin-treated (metformin+) T2D cases compared to metformin-negative (metformin-) T2D or ND controls (*P*-values from Wilcoxon test). **(d)** SIAMCAT models can easily distinguish between metformin+ T2D and ND controls and between metformin+ T2D cases and metformin- T2D cases. On the other hand, metformin- T2D cases and ND controls are harder to distinguish. See Figure 1b in Forslund *et al.* as reference.

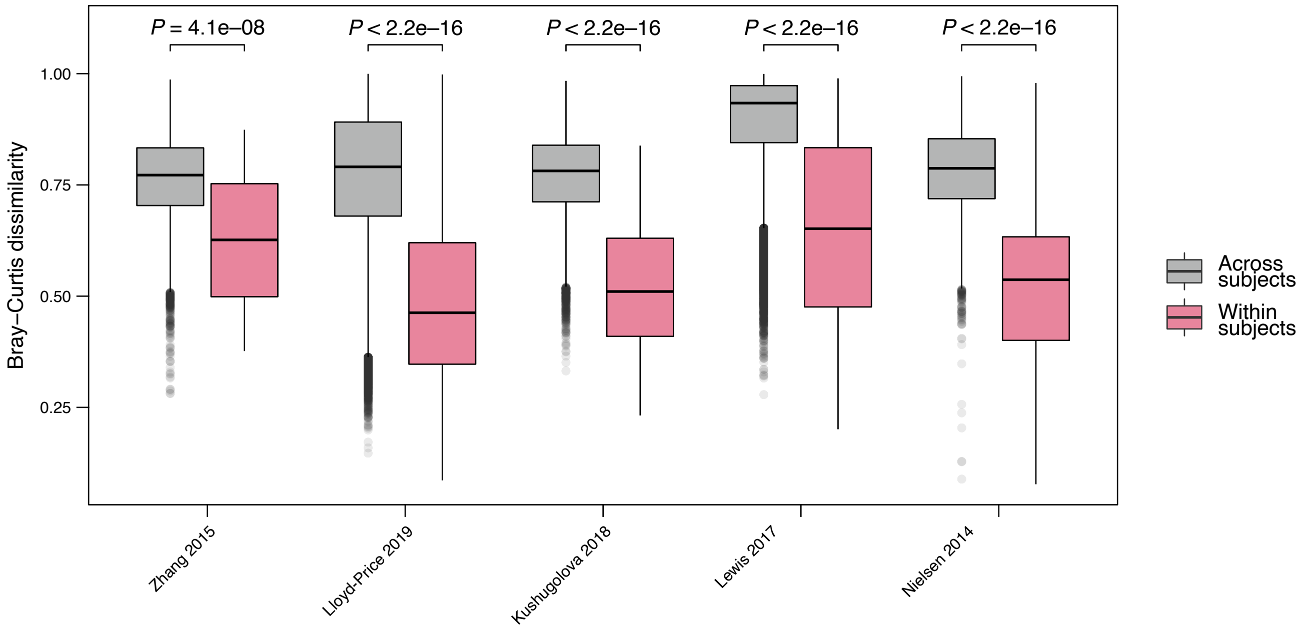


**Supplementary Figure 3: Metagenomic samples are more similar within subjects than across subjects**
To illustrate that metagenomic measurements are more similar within subjects than across subjects, all pairwise Bray-Curtis dissimilarities were calculated for in-house datasets with repeated measurements for different subjects. Dissimilarity values are displayed as boxplots and coloured depending on whether the two samples came from the same subject or from two different subjects. The dissimilarities for samples from the same subject are significantly lower than across subject for all datasets (*P*-values from Wilcoxon test). Boxes denote the IQR across all values with the median as a thick black line and the whiskers extending up to the most extreme points within 1.5-fold IQR.

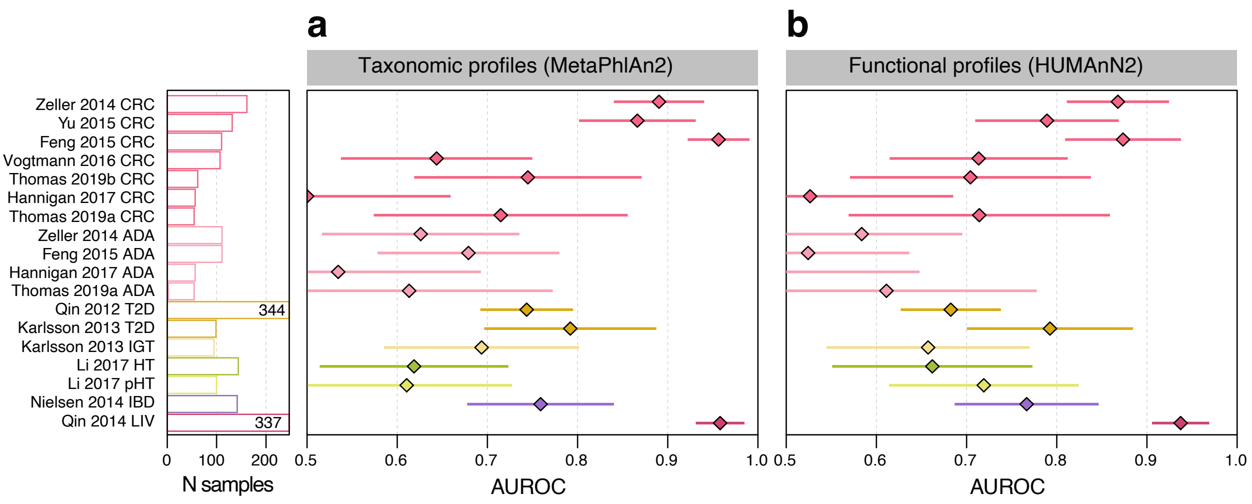


**Supplementary Figure 4: Large-scale application of the SIAMCAT machine learning workflow to human gut metagenomic disease association studies in the curatedMetagenomicsData package.**
**(a)** Application of SIAMCAT machine learning workflows to taxonomic profiles generated from fecal shotgun metagenomes using MetaPhlAn2 as available from *curatedMetagenomicData* (Pasolli *et al.* 2017). Cross-validation performance for discriminating between diseased patients and controls quantified by the area under the ROC curve (AUROC) is indicated by diamonds (95% confidence intervals denoted by horizontal lines) with sample size per dataset given as additional panel (cut at N = 250 and given by numbers instead). See **Table 1** and **Supplementary Table 1** for information about the included datasets and key for disease abbreviations. **(b)** Application of SIAMCAT machine learning workflows to functional profiles obtained from HUMAnN2 as provided by *curatedMetagenomicData* (Pasolli *et al.* 2017) for the same datasets as in (a).

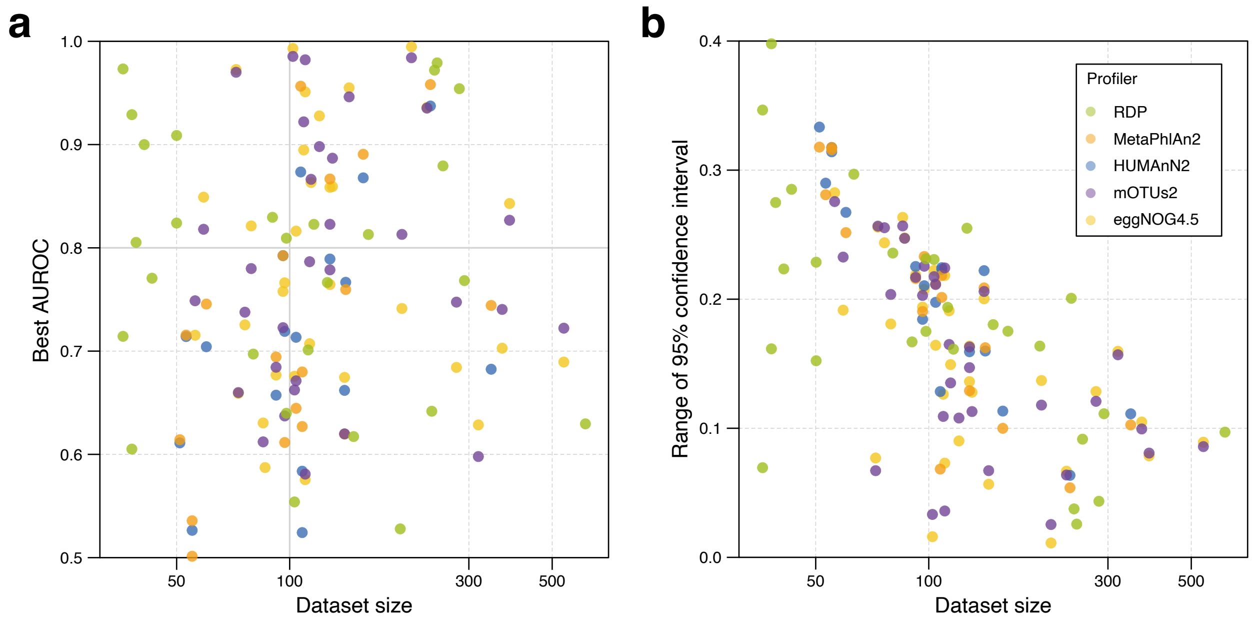


**Supplementary Figure 5: Dataset size relates to classification accuracy and the AUROC confidence interval**

**(a)** Dataset size is plotted against model accuracy as measured by AUROC for the best-performing parameter set (see **Figure 4** in the main text and **Supplementary Figure 4**). Dots are colored by the input data type. 57% of the classification tasks based on datasets with 100 or more samples could be classified with an AUROC of 0.75 or higher compared to only 35% of classification tasks based on datasets with fewer than 100 samples. **(b)** Dataset size is plotted against the range of the 95% confidence interval for the best AUROC. Dots are colored by the input data type. There is a clear trend toward an overall lower range of the confidence interval for bigger datasets.

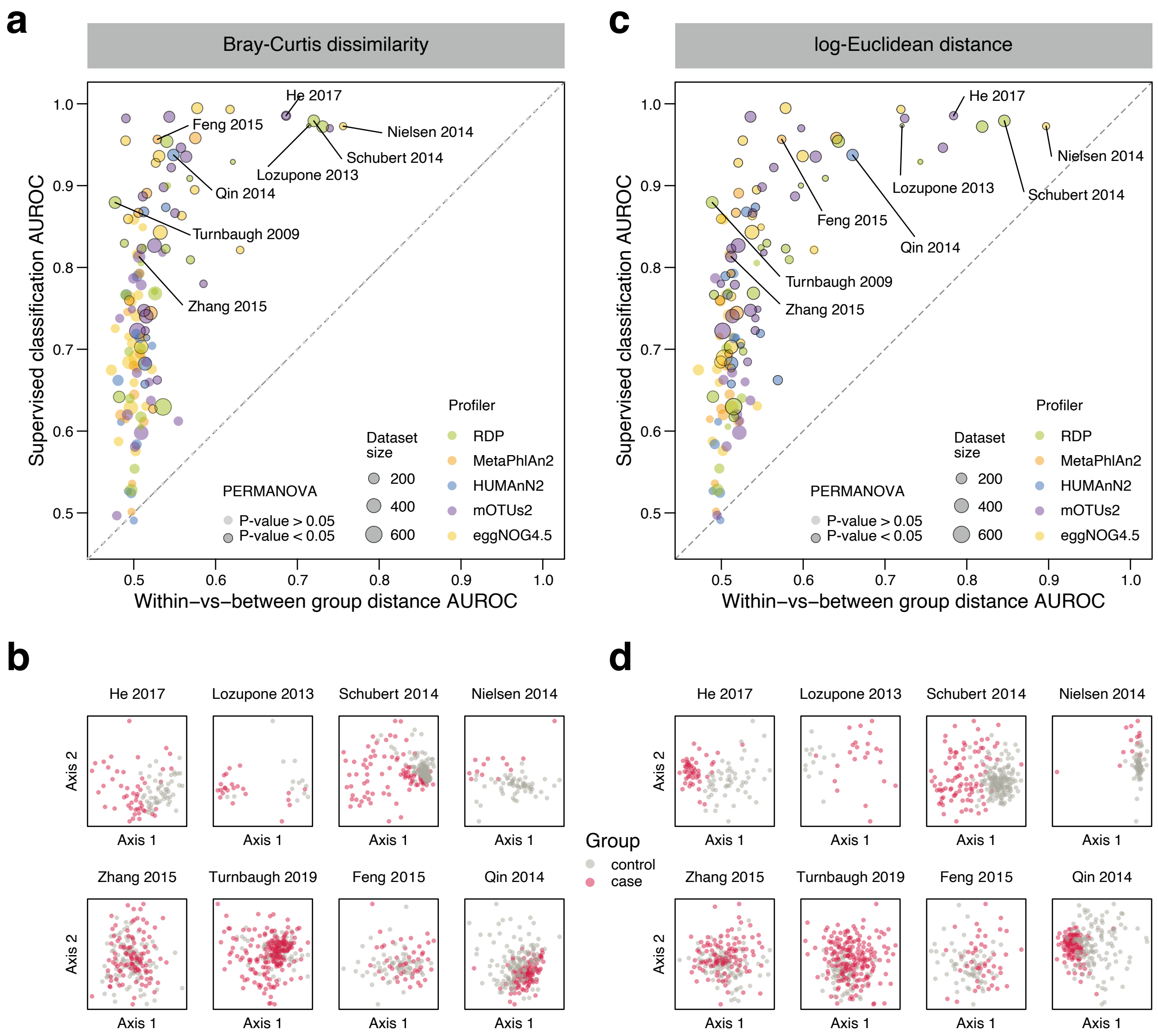


**Supplementary Figure 6: Machine learning can distinguish group differences even when samples can not be separated based on common ecological distances**

**(a)** Machine learning model accuracy as measured by AUROC for the best-performing parameter set (see **Figure 4** in the main text and **Supplementary Figure 4**) is plotted against the AUROC quantifying the separation of groups on the Bray-Curtis distance (see Methods). Dot size is proportional to the number of samples for each classification task and dots are colored by data type. Dot outline indicates if the *P*-values calculated by PERMANOVA is below 0.05 or above. For many classification tasks, our analysis indicates no separation of groups based on the Bray-Curtis distance, whereas machine learning models can be trained to accurately distinguish between the two groups. **(b)** Principal Coordinate plots based on the Bray-Curtis distance for the classification tasks highlighted in (a). Control samples are shown as grey dots and disease samples are shown in red irrespective of the disease. For the classification tasks in the upper row, there is a good separation both based on the Bray-Curtis distance and the machine learning analysis. For the tasks in the lower row, however, accurate machine learning models can be trained but there is no apparent separation based on the Bray-Curtis distance. **(c)** Equivalent plot as in (a), but based on the Euclidean distance after log-transformation. **(d)** Equivalent plots as in (b), but based on the Euclidean distance after log-transformation.

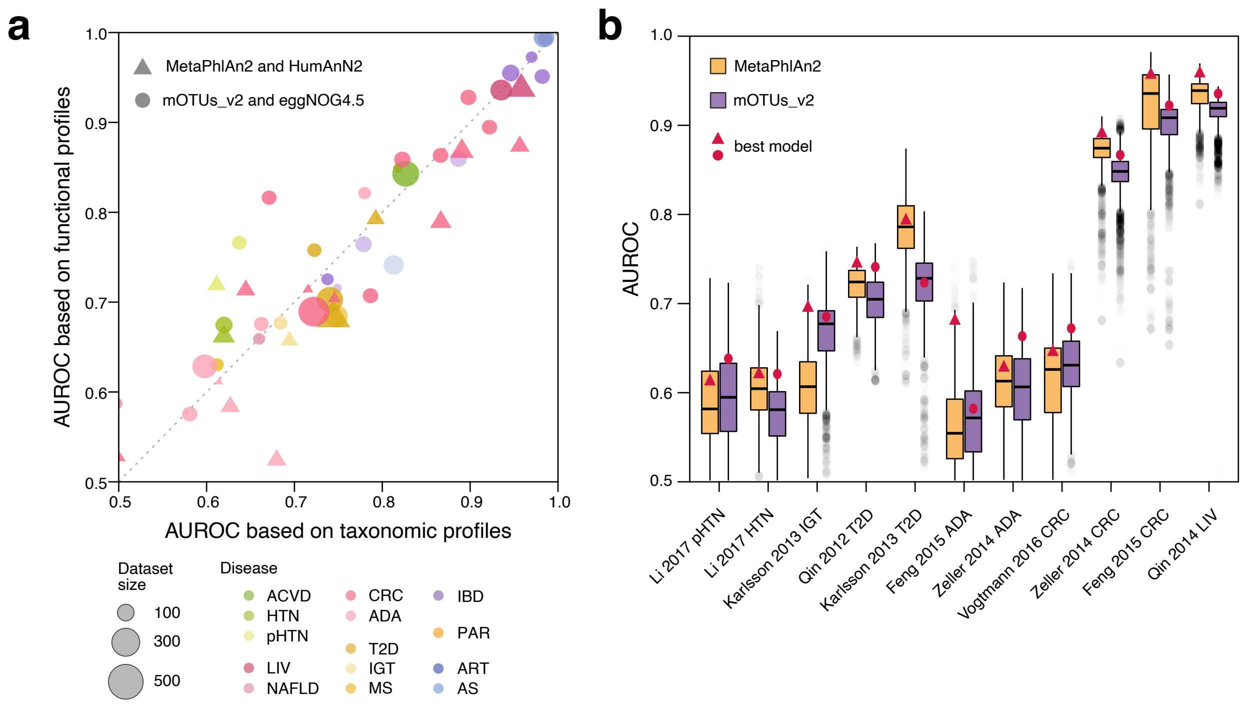


**Supplementary Figure 7: Classification accuracy is not impacted by choice of profiler**
**(a)** Model accuracy as measured by AUROC is plotted for the best-performing parameter set (see **Figure 4** in the main text and **Supplementary Figure 4**) for taxonomic and functional profiles derived from the same dataset. Profiles from mOTUs2 and eggNOG4.5 are plotted against each other as are profiles from MetaPhlAn2 and HUMAnN2. Overall, the accuracies are very well correlated (Pearson's r = 0.91, *P* < 2e-16), indicating that taxonomic and functional profiles lead to very similar model performances across a wide range of classification tasks. Dot size is proportional to the number of samples per classification task and dots are colored according to the disease. See **Table 1** for a key of the disease abbreviations. **(b)** For those classification tasks that involve the same dataset and the same disease and for which both mOTUs2 and MetaPhlAn2 profiles are available, all AUROC values from the complete parameter set exploration are shown as boxplots with the color indicating the two different profilers. The AUROC values for the best-performing parameter set (see **Figure 4** in the main text and **Supplementary Figure 4**) are indicated by dots and triangles, respectively. Although there are differences between mOTUs2 and MetaPhlAn2 on individual datasets, there is no clear trend towards either method, indicating that the choice of taxonomic profiler does not significantly impact the resulting model accuracy (*P* = 0.41 from paired Wilcoxon test with the best-performing AUROC values). Boxes denote the IQR across all values with the median as a thick black line and the whiskers extending up to the most extreme points within 1.5-fold IQR.

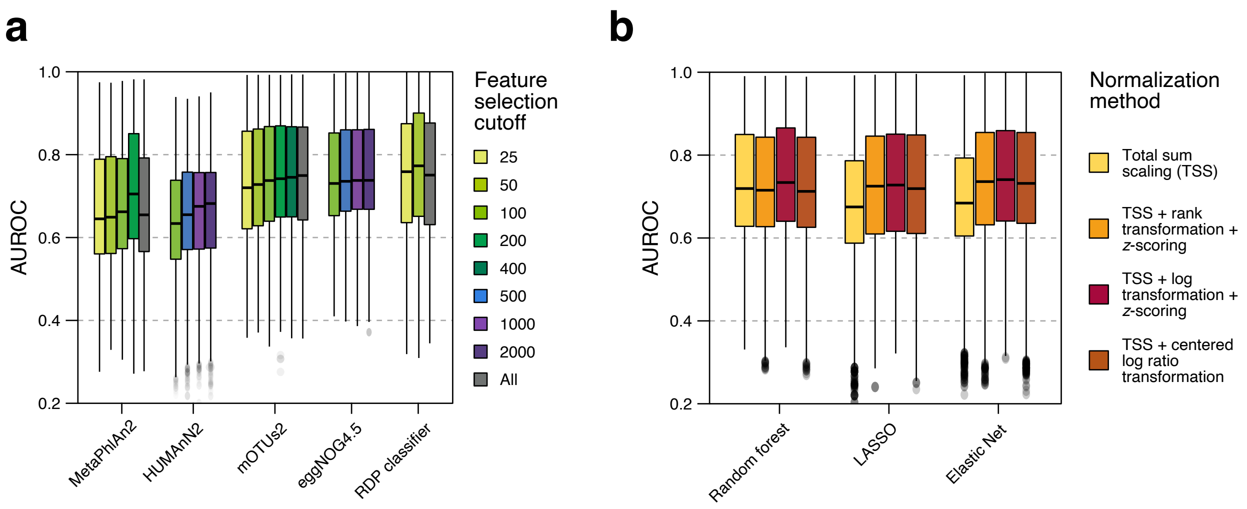


**Supplementary Figure 8: Influence of feature selection cutoff and normalization method on classification accuracy**

**(a)** Accuracy as measured by AUROC is displayed for all parameter combinations and all datasets of different input types broken down by different cutoffs in the feature selection procedure. "All" indicates that the feature selection was turned off and all possible features were used for training of the machine learning model. Whereas the choice of feature selection cutoff is less important for profiles generated with the RDP profiler, the other data types seem to profit the more features are included, especially in the case of HUMAnN2 profiles. Boxes denote the IQR across all values with the median as a thick black line and the whiskers extending up to the most extreme points within 1.5-fold IQR. **(b)** Accuracy as measured by AUROC is displayed for all parameter combinations and all datasets of different input types broken down by the different normalization methods included in the parameter exploration. The resulting accuracy is barely impacted by the choice of normalization method when the model is trained with the Random Forest classifier. For the other two machine learning algorithms, however, the naive total sum scaling (TSS) normalization is not sufficient for optimal performance. Boxes denote the IQR across all values with the median as a thick black line and the whiskers extending up to the most extreme points within 1.5-fold IQR.

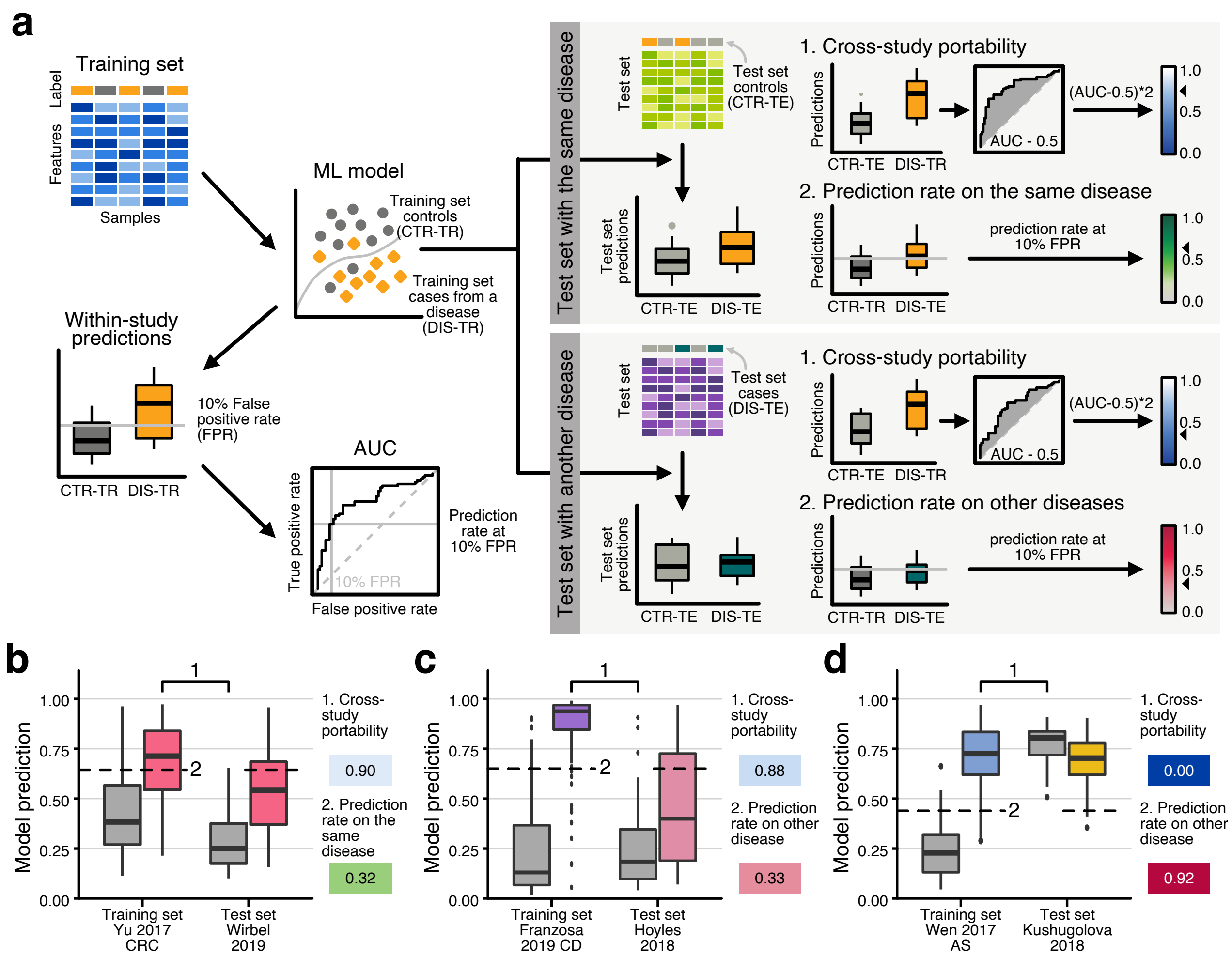


**Supplementary Figure 9: Measures to evaluate the cross-study application of ML models**
**(a)** Given a training dataset containing controls (CTR-TR) and cases for a specific disease (DIS-TR), a ML model can be trained with SIAMCAT to distinguish between cases and controls. This ML model then produces within-study predictions (on samples that were left out during cross-validation), on the basis of which model performance is calculated as AUC (area under the ROC curve). Additionally, we determine the decision boundary that corresponds to a 10% false positive rate (FPR), that is, at which cutoff for the model predictions would 10% of the control samples be incorrectly classified as diseased.
When the trained ML model is applied to samples from other datasets, two situations can arise: either the test set contains controls (CTR-TE) and cases (DIS-TE) from the same disease as the test set (top box) or it contains CTR-TE and DIS-TE from another disease (bottom box). In either case, predictions made by applying the ML model to the test set are used as a basis for evaluating model performance in cross-study transfer. This is assessed by two different measures.
First, we calculate cross-study portability from the AUC between CTR-TE and DIS-TR (by scaling it to the interval between 0 and 1). This measure captures how well external controls, CTR-TE, can be separated from the cases contained in the cross-validation data set, DIS-TR (analogous to the AUC-based performance evaluation within the cross-validation set). Low cross-study portability values indicate that there is no separation between cases and external controls, meaning that the model would show and increased false-positive rate on control samples from other datasets.
Second, we calculate the prediction rate for external cases, DIS-TE, at a prediction cutoff that corresponds to a 10% FPR on the cross-validation data set. This measure quantifies to which extent the model exhibits an elevated false positive rate for other diseases, which could be due to technical differences between studies (which would also be reflected in a low cross-study portability) or due to biological similarity between diseases (if the same microbial markers are enriched in both diseases, one would expect an elevated prediction on the other disease as well). This measure is thus a proxy for the disease-specificity of the ML model. For data sets with cases from the same disease, this evaluation amounts to assessing prediction rate (of the same disease) across data sets.
**(b)**, **(c)**, and **(d)** illustrate the within-study and test set predictions for selected examples from our ML meta-analysis for transfer across data sets for the same (a) or different diseases (c), (d) with (d) presenting an extreme example of issues with both cross-study portability and disease-specificity. Numbers indicate the type of comparisons performed across data sets (as indicated on the right-hand side of the plots).

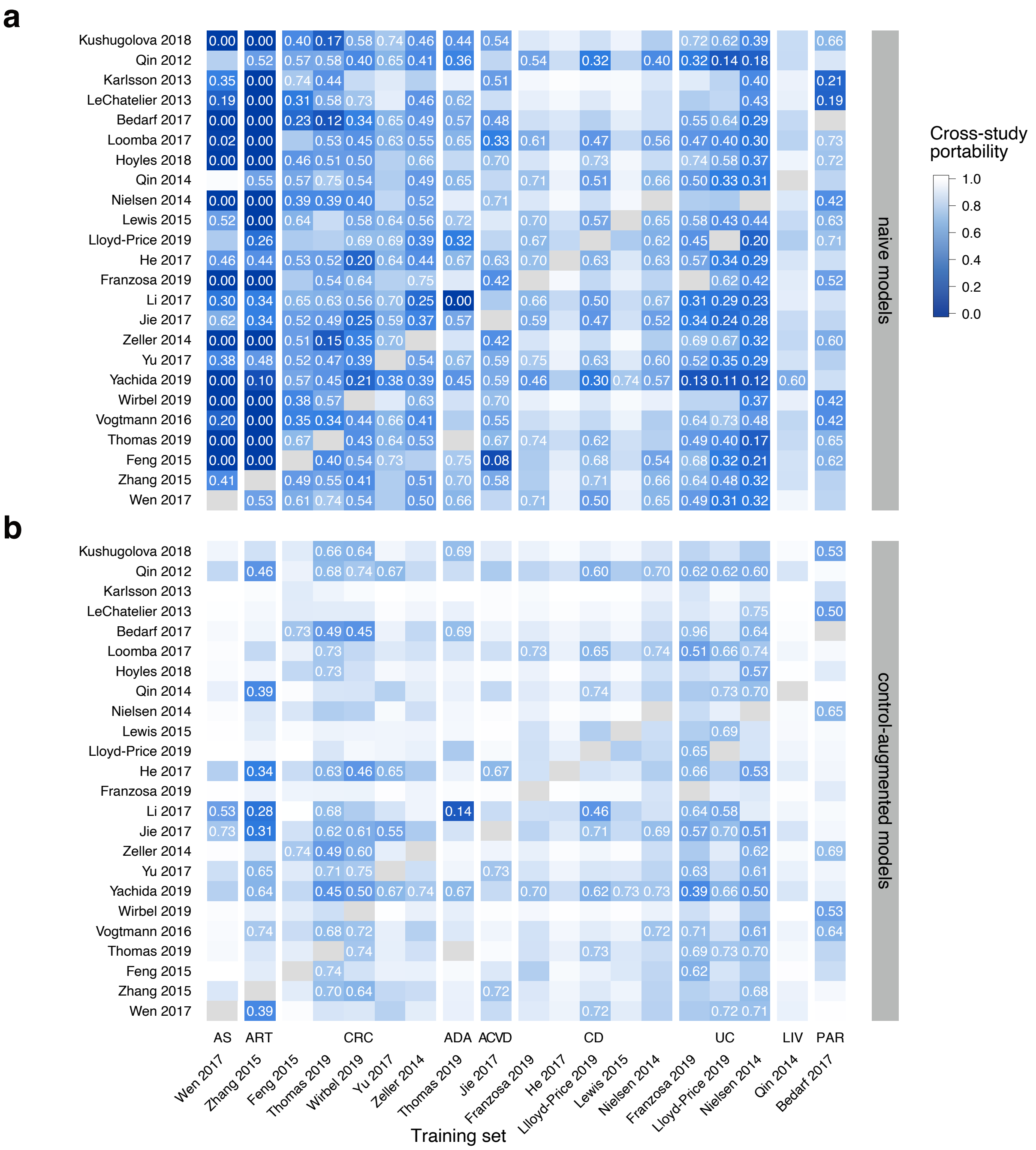


**Supplementary Figure 10: Naive ML models show lower cross-study portability when applied to external datasets compared to control-augmented models**

Cross-study portability on the control portion of external studies (see Methods) is shown as a heatmap for naive models **(a)** and control-augmented models **(b)**. The heatmap only includes models with an AUROC of 0.75 or higher (see **Figure 5** in the main text). Values equal or smaller than 0.75 are highlighted.

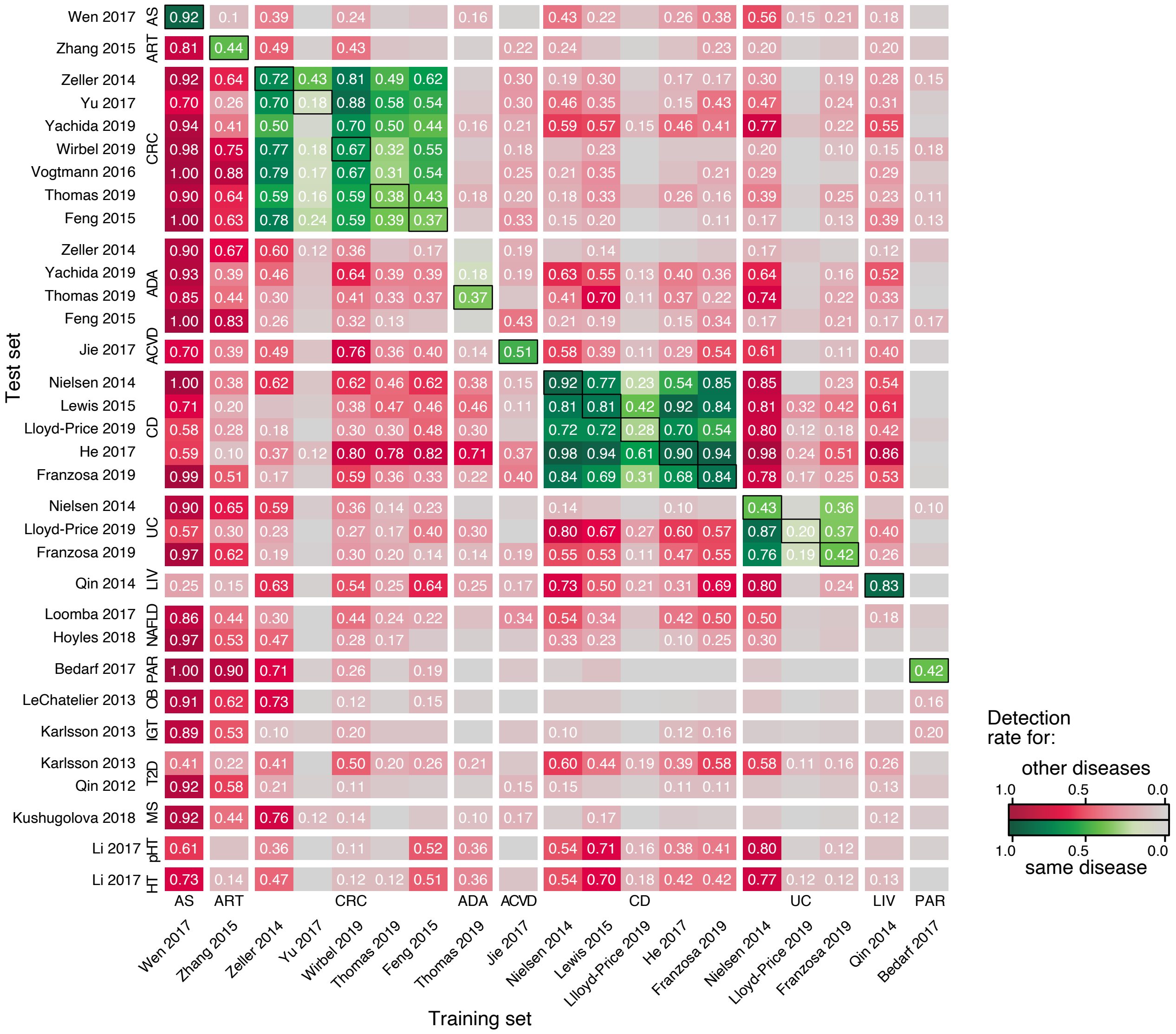


**Supplementary Figure 11: Naive ML models make a high level of false predictions on external datasets** Detection rates for other diseases (red color-scheme) or the same disease (green color-scheme) are shown for the naive ML models. True positive rates of the models, when applied to the training set in cross-validation, are indicated by boxes around the tile. Detection rates over 10% are labeled. When compared to **Figure 5d** in the main text, the naive ML models show dramatically higher detection rates on other diseases compared to control-augmented ML models.

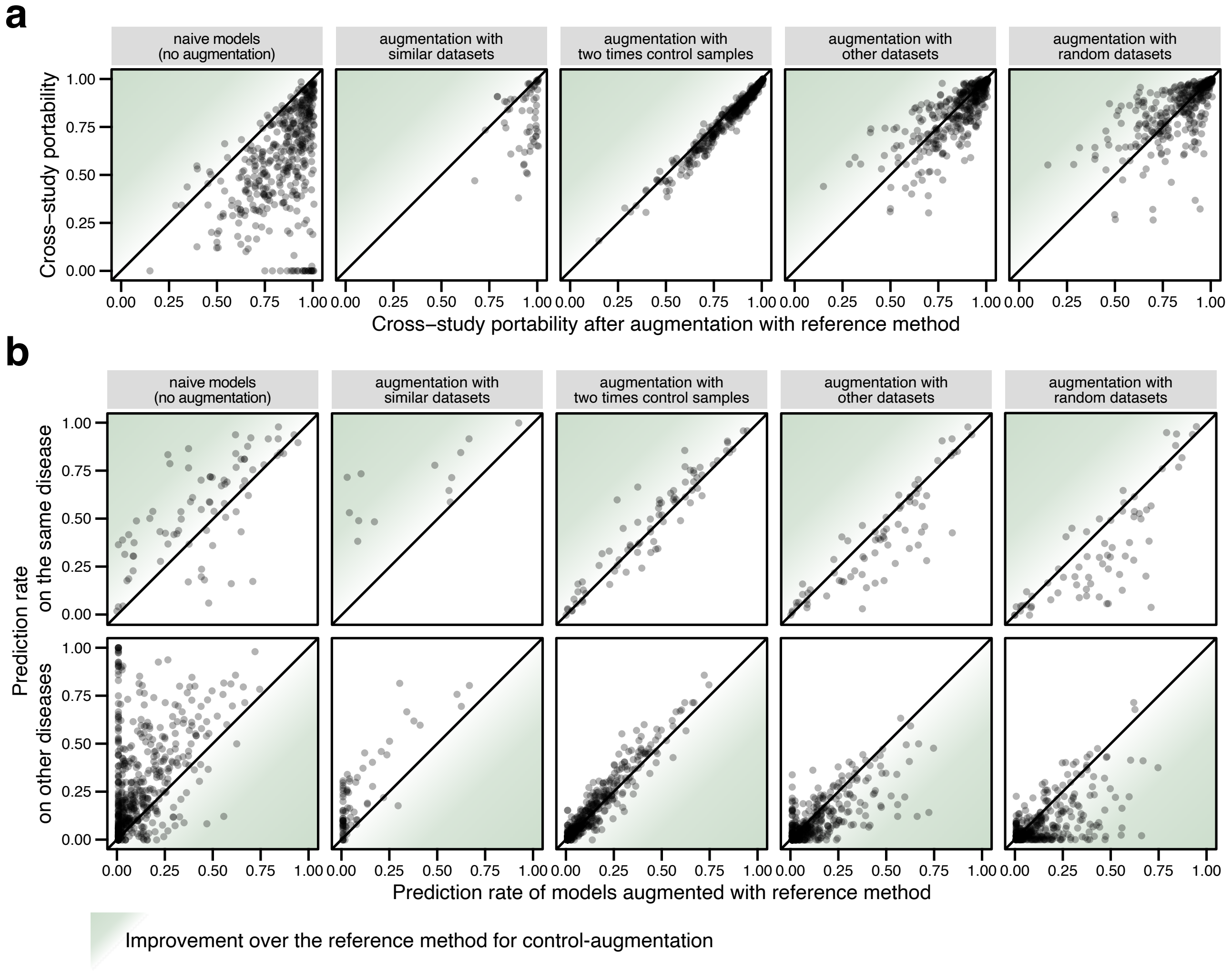


**Supplementary Figure 12: Control-augmentation strategy generally improves model transfer without a strong dependence on the type and number of controls sampled**

**(a)** Cross-study portability and **(b)** prediction rate on the same disease (upper row) and on other diseases (lower row) are compared for all models between the reference method for control augmentation (as described and displayed in the main text) and variations of that approach. The reference method consisted in the addition of five times the number of controls sampled randomly in each cross-validation split from a set of three large (>250 samples) cohort studies (see Methods in the main text). The other approaches are defined as follows:

The **naive models** are models without augmentation, shown as a baseline to visualize the improvements in model transfer achieved by control-augmentation (reference method). While cross-study portability and disease-specificity generally improve, the prediction rate on the same disease (e.g. for different studies including cases of colorectal cancer or Crohn's disease) is sometimes reduced by control-augmentation.

**Control-augmentation with similar datasets** was used for a subset of datasets in our ML meta-analysis that clustered together (all datasets included samples from the same population and were generated in the same laboratory; Yu et al. *Gut* 2017, Jie et al. *Nat comun* 2017, He et al. *Gigascience* 2017, and Qin et al. *Nature* 2012). When training on those datasets, controls were randomly sampled from the other datasets listed above. Using similar datasets in the control-augmentation does not lead to the same improvements that are seen when using a more diverse set of studies to augment.

**Control-augmentation with two times the number of control samples** is very similar to the reference method in that controls are sampled from the same cohort studies as in the reference method but a lower number of controls is added (twice the number instead five times the number of controls). The method performs very similarly albeit slightly worse compared to the reference method.

For the **control-augmentation with other datasets**, we sampled twice the number of control samples from another pool of datasets (Danish samples from Nielsen et al. *Nat Biotech* 2014, samples from mothers in Backhed et al. *Cell Host & Microbe* 2015, Vincent et al. *Microbiome* 2016, Zhu et al. *Microbiome* 2018, and Poyet et al. *Nat Med* 2019); more data sets were necessary to obtain a sufficient number of control samples for the reference control-augmentation approach. We used only samples reported as controls and filtered out repeated samples from the same subject, whenever applicable. The results of this methods are also similar to the reference approach, but lead to a further decrease in prediction on the same disease.

Lastly, we used **control-augmentation with random datasets**, for which we randomly sampled five datasets out of the meta-analysis set and used their control samples to augment the training set. The resulting augmented model was not evaluated for model transfer on the datasets which were used for augmentation. This method behaves similarly to the control-augmentation with other datasets with only minor differences to the reference method.

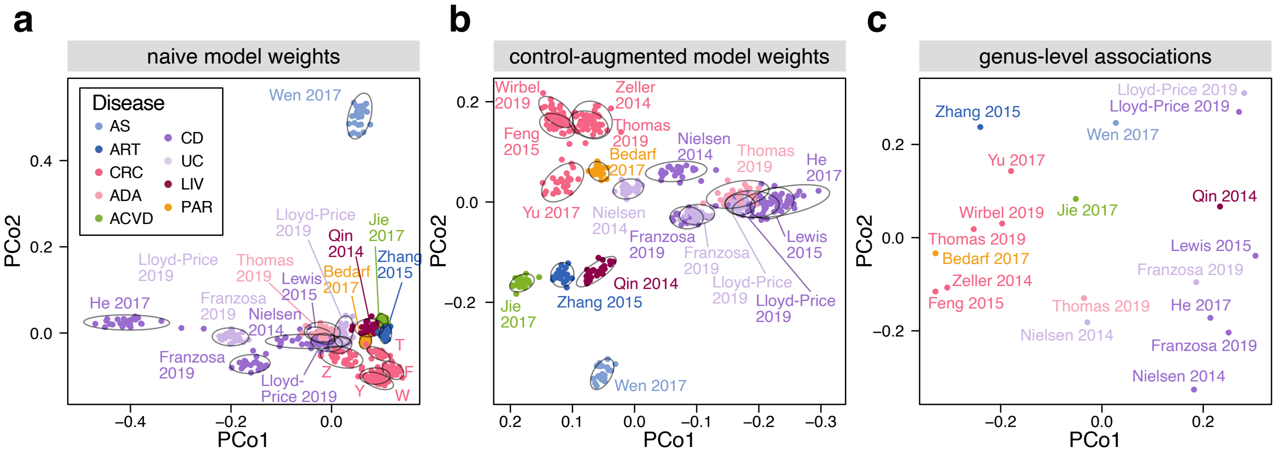


**Supplementary Figure 13: Datasets cluster by disease, both when considering ML model weights or associations**

**(a)** Principal coordinate (PCo) analysis based on Canberra distances between relative models weights for naive ML models. Each dot represents a trained model from the repeated cross-validation. Datasets are indicated by 90% density ellipses. For more convenient labeling, the CRC datasets are abbreviated by their first letter. **(b)** PCo analysis based on Canberra distances between relative models weights for control-augmented ML models. Each dot represents a trained model from the repeated cross-validation and datasets are again indicated by 90% density ellipses. **(c)** PCo analysis based on the Canberra distances between genus-level generalized fold changes for each dataset.

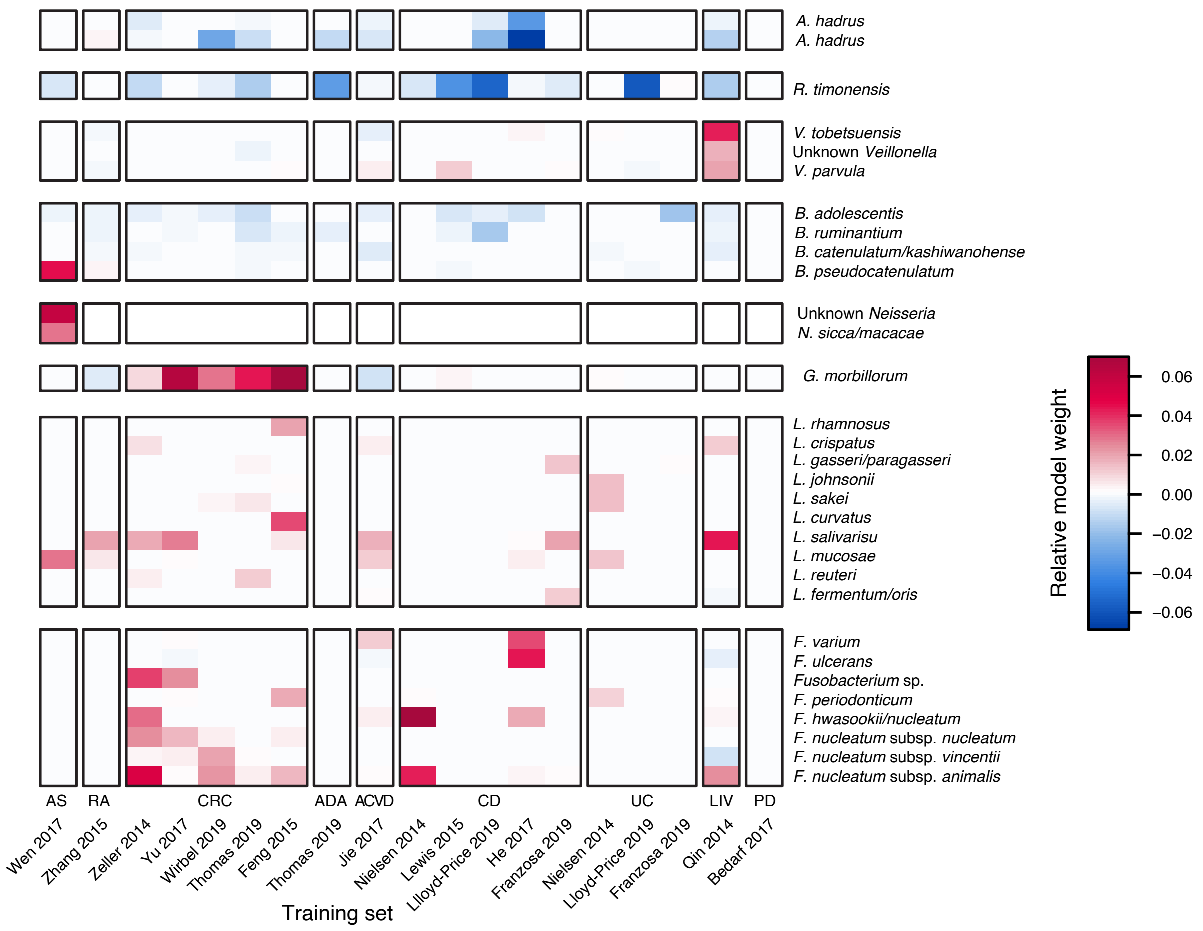


**Supplementary Figure 14: ML model weights reveal shared and disease-specific predictors**
Heatmap showing the relative weights of control-augmented ML models for mOTUs in selected genera. Blue values indicate that the mOTU is a control-enriched predictor and red values indicate a disease-enriched predictor.

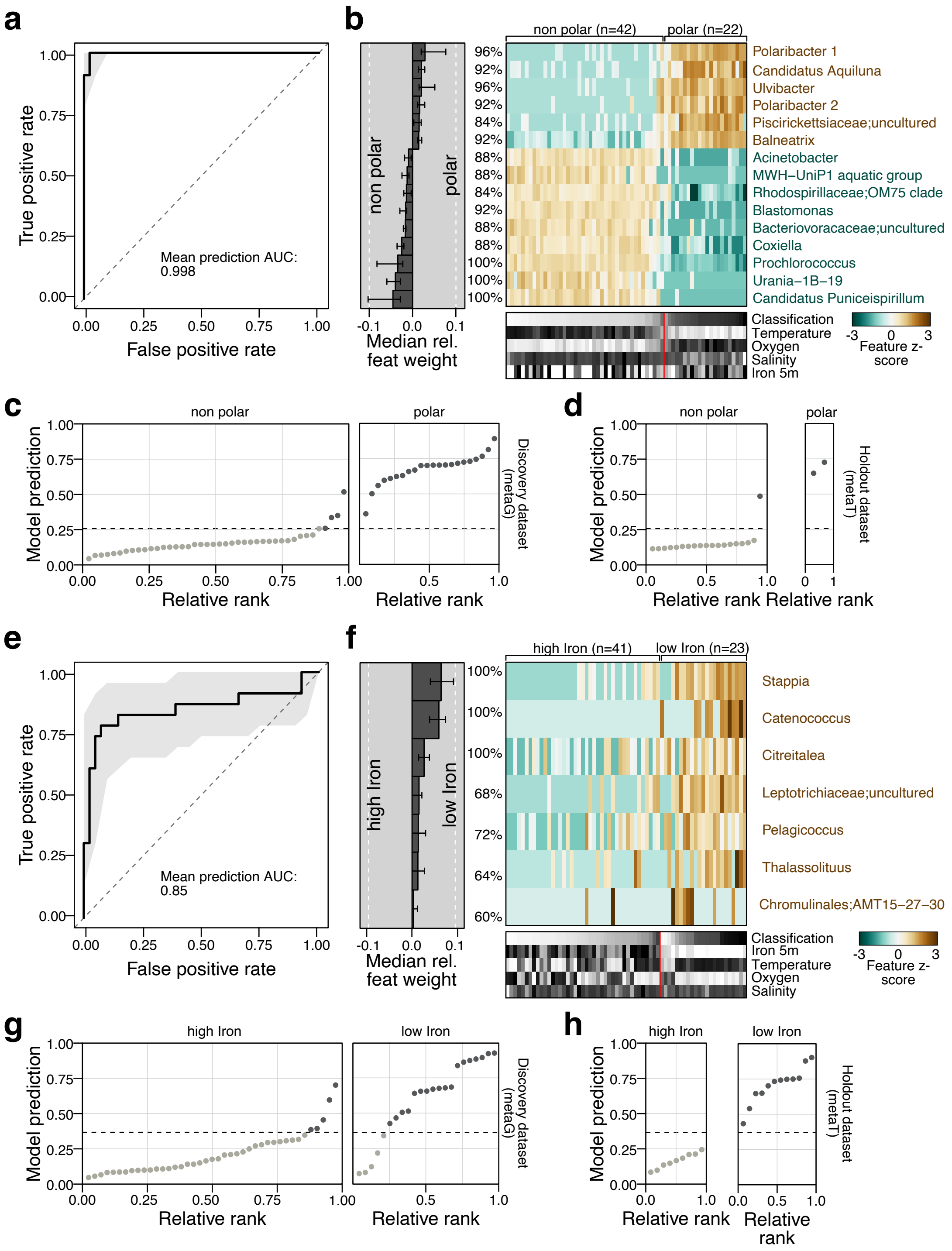


**Supplementary Figure 15: SIAMCAT can be applied to metagenomic and metatranscriptomic measurements from environmental samples**

**(a)** A LASSO model trained with SIAMCAT can distinguish samples from polar ocean environments from non-polar ocean samples with an AUROC of 0.998. **(b)** The model interpretation plot generated by SIAMCAT shows the distribution across all samples for those genera which are most predictive of polar and non-polar ocean environments (central heatmap). The importance of these genera in the model are shown as barplot on the left. Below the central heatmap, other environmental measurements are shown together with the model predictions. **(c)** Ranked model predictions are plotted for all metagenomic ocean samples used for training the model, separated by polar and non-polar ocean environments. The dotted line represents the cutoff for the predictions that corresponds to a false positive rate of 10%. **(d)** Dots show the ranked model predictions derived from applying the trained model to meta-transcriptomic ocean samples, separated by polar and non-polar ocean environments. The dotted line represents the cutoff for the predictions that corresponds to a false positive rate of 10% on the metagenomic discovery set (see **(c)**). **(e)** A LASSO model trained with SIAMCAT can distinguish samples from low-iron ocean environments from high-iron ocean samples with an AUROC of 0.85. **(f)** The model interpretation plot generated by SIAMCAT shows the distribution across all samples for those genera which are most predictive of high and low-iron ocean environments (central heatmap). The importance of these genera in the model are shown as barplot on the left. Below the central heatmap, other environmental measurements are shown together with the model predictions. **(g)** Ranked model predictions are plotted for all metagenomic ocean samples used for training the model, separated by high and low-iron ocean environments. The dotted line represents the cutoff for the predictions that corresponds to a false positive rate of 10%. **(h)** Dots show the ranked model predictions derived from applying the trained model to meta-transcriptomic ocean samples, separated by high and low-iron ocean environments. The dotted line represents the cutoff for the predictions that corresponds to a false positive rate of 10% on the metagenomic discovery set (see **(g)**).
